# Mitogenomic differentiation in spinner (*Stenella longirostris*) and pantropical spotted dolphins (*S. attenuata*) from the eastern tropical Pacific Ocean

**DOI:** 10.1101/091215

**Authors:** Leslie Matthew S., Archer Frederick I., Morin Phillip A.

## Abstract

Spinner dolphins (*Stenella longirostris*) and spotted dolphins (*S. attenuata*) show high intraspecific morphological diversity and endemic subspecies in the eastern tropical Pacific Ocean (ETP). Previous studies of mitochondrial DNA (mtDNA) have found low genetic differentiation among most of these groups, possibly due to demographic factors, ongoing gene flow, and/or recent divergence. These species were heavily depleted due to bycatch in the ETP yellowfin tuna fishery. Because population structure is important for accurate management of the recovery of these species, we collected whole mitochondrial genome sequences from 104 spinner and 76 spotted dolphins to test structure hypotheses at multiple hierarchical levels. Our results showed significant differences between subspecies of spotted (*F*_ST_: 0.0125; *P =* 0.0402) and spinner dolphins (*F*_ST_: 0.0133; *P* = 0.034), but no support for the division of existing offshore stocks of spotted dolphins or Tres Marias spinner dolphins. We compare these results to previous results of genome-wide nuclear SNP data and suggest high haplotype diversity, female dispersal, male philopatry, or relative power of the two datasets explains the differences observed. Our results further support a genetic basis for biologically meaningful management units at the subspecies level, and provide a critical component to mitigating historical and continued fishery interactions.

## Introduction

Determining population genetic structure is important for accurately managing protected wildlife species (Taylor 2005). Because mitochondrial DNA (mtDNA) is more abundant in cells and has a higher rate of mutation - thus accruing variability on a time-scale typical of population divergence – it has been the preferred marker for population genetic studies of wildlife (Moritz 1994; Allendorf 2017). Moreover, because of the strictly maternal inheritance of mtDNA, comparing the strength of genetic structure between mtDNA and nuclear DNA (nuDNA) can provide valuable insights into maternal genetic structure and sex-bias dispersal in wildlife populations (Moritz 1994). Mitochondrial DNA data are particularly useful for species with strong matrilineal social structure – such as several toothed whale species (i.e., killer whales, sperm whales, and pilot whales). In cetaceans, whole mtDNA genome (mitogenome) sequencing has provided additional clarity species-level population structure and phylogeographic patterns where single mtDNA markers have not (Archer *et al.* 2013; Morin *et al.* 2010). We expanded upon previous mtDNA datasets and include the whole mitochondrial genome to test for population structure in two species of dolphin.

Fisheries bycatch is arguably the largest threat facing cetaceans today (Read *et al.* 2006). One of the largest fisheries bycatch events in history occurred in the eastern tropical Pacific Ocean (ETP) and heavily impact pelagic spinner and spotted dolphins in this area. These two species were abundant (numbering in the low millions) in the ETP (Wade *et al.* 2007), but because both species commonly associate with one-another and with large tuna (see Scott *et al.* 2012 for details), bycatch in the dolphin-set tuna purse-seine fishery starting in the 1960s killed hundreds of thousands annually (Lo and Smith 1986, National Research Council 1992, Wade 1995). Despite protection under the U.S. Marine Mammal Protection Act of 1972 and multi-national protection under the 1999 Agreement on the International Dolphin Conservation Program (Joseph 1994, Gosliner 1999), ETP spinner and spotted dolphin population abundances remain low (Wade *et al.* 2007, Gerrodette *et al.* 2008). Determining how populations are naturally structured in the ETP is critical to accurately managing the recovery of these populations.

## ETP Spinner Dolphins

Globally, there are four subspecies of spinner dolphin (*Stenella longirostris*). The nominate form, the pantropical spinner (*S. l. longirostris*) inhabit in all tropical waters of the world outside the ETP. Pantropical spinners are usually associated with islands in the central and western Pacific, such as the Hawai‘ian Islands. In shallow waters of Southeast Asia, there is a much smaller dwarf spinner subspecies (*S. l. roseiventris*) (Perrin *et al.* 1989, 1999). The Central American spinner dolphin (*S. l. centroamericana*) and the eastern spinner dolphin (*Stenella l. orientalis*) are endemic to the ETP (Fig. 1, based on Perrin 1985). Analyses of external coloration, body size, and the extensive analyses of cranial morphology lead to the erection of these ETP subspecies (Perrin *et al.* 1991, Douglas *et al.* 1992). The Central American subspecies is found off the Pacific coasts of Southern Mexico south through Panama, in relatively near-shore waters. The eastern spinner dolphin (*S. l. orientalis*), on the other hand, inhabits offshore waters that extend from Baja California, Mexico, south to Ecuador (Perrin 1990).

**Figure 1.**
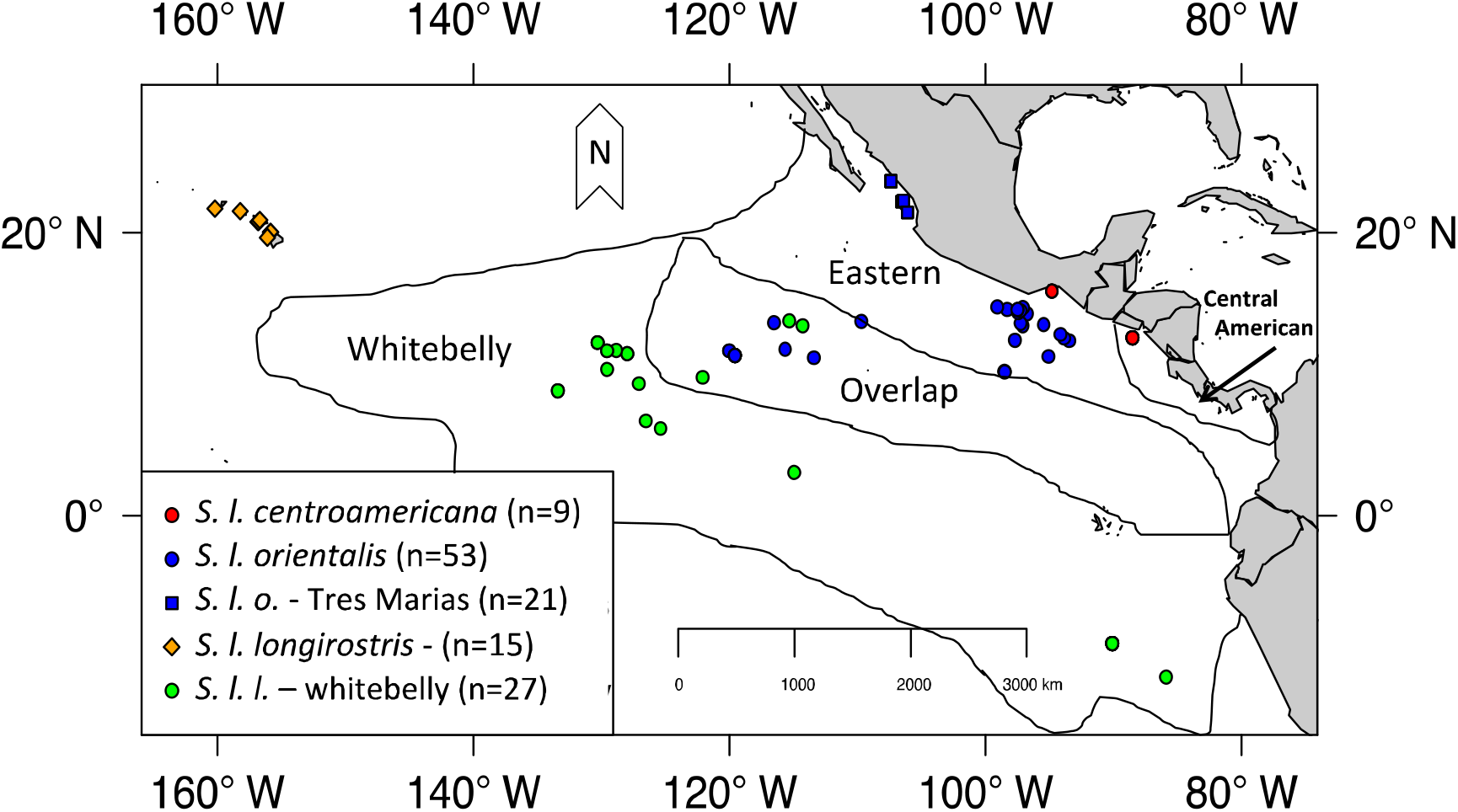
Sampling localities and range map for spinner dolphins within the ETP. Subspecies and stock boundaries based on Perrin *et al*. 1985. Red dots indicate Central American spinners. Blue symbols indicate eastern spinners - boxes are the proposed Tres Marias form. Green dots indicate whitebelly spinners, a proposed intergrade between the pantropical (orange diamonds) and the eastern subspecies. mtDNA sample sizes are in the legend.

For management purposes, the two ETP endemic spinner dolphin subspecies are considered stocks, plus a third stock - the whitebelly spinner. The “whitebelly” spinner is proposed to represent a hybrid swarm between the eastern subspecies and the pantropical subspecies of the central and western Pacific (Perrin *et al.* 1991). Taxonomically, it is classified as part of the nominate (pantropical) spinner subspecies *S. l. longirostris*. Significant geographic overlap exists between the eastern subspecies and the whitebelly form (Perrin *et al.* 1985) (See Fig. 1). Finally, a distinct morphotype of the eastern spinner dolphin, known as the “Tres Marias” spinner dolphin, has been described from near the islands of the same name off the coast of Mexico. These were thought to be a distinct type based on external body morphometrics (Perryman and Westlake 1998).

Some molecular genetics approaches have not found genetic structure corresponding to the subspecific morphological differences (Dizon *et al.* 1994, Galver 2002). Andrews *et al.* (2013) estimated high levels of gene flow between subspecies in the ETP using autosomal and mitochondrial genes and found a shared Y chromosome haplotype in the eastern and Central American subspecies that was not found in the pantropical or dwarf subspecies. Interestingly, this locus was found to be polymorphic in whitebellies, supporting the hypothesis of introgression in this form (Andrews *et al.* 2013). The authors proposed that sexual selection was driving the divergence of spinner dolphins in the ETP. Recently, Leslie and Morin (2016) found strong population structure within both species using genome-wide SNP data.

## ETP Spotted Dolphins

The pantropical spotted dolphins (*Stenella attenuata*) in the ETP is split into two subspecies based on morphometric analyses: a coastal endemic subspecies (*S. a. graffmani -* Perrin 1975, Perrin *et al.* 1987) and an offshore pantropical subspecies (*S. a. attenuate*). Genetic analyses of microsatellites show high genetic diversity in spotted dolphins and support some differentiation between subspecies (Escorza-Treviño *et al.* 2005). This study identified at least four demographically independent populations within the coastal subspecies (*S. a. graffmani*) and differences between southern populations of the coastal subspecies and the pelagic subspecies. However, they found no differences between the northern populations of the coastal subspecies and the pelagic subspecies. Escorza-Treveño *et al.* (2005) identified demographically independent populations within the coastal subspecies and posited that interchange continues between the northern *S. a. graffmani* populations and the offshore pantropical subspecies.

Although the results of Escorza-Treviño *et al.* (2005) indicate substructure, the entire coastal subspecies is currently a single management stock. Offshore pantropical spotted dolphins in the ETP are divided into two stocks: 1) the ‘northeastern’ (NE) stock is defined geographically as north of 5°N, east of 120°W, and 2) the ‘western-southern’ (WS) stock is defined as south and west of this northeastern area (Fig. 2) (Perrin *et al.* 1994). A distributional hiatus along 5°N is the basis for the north-south boundary between NE and WS stocks (Perrin *et al.* 1994), and this has recently been supported by SNP analyses (Leslie and Morin 2016).

**Figure 2.**
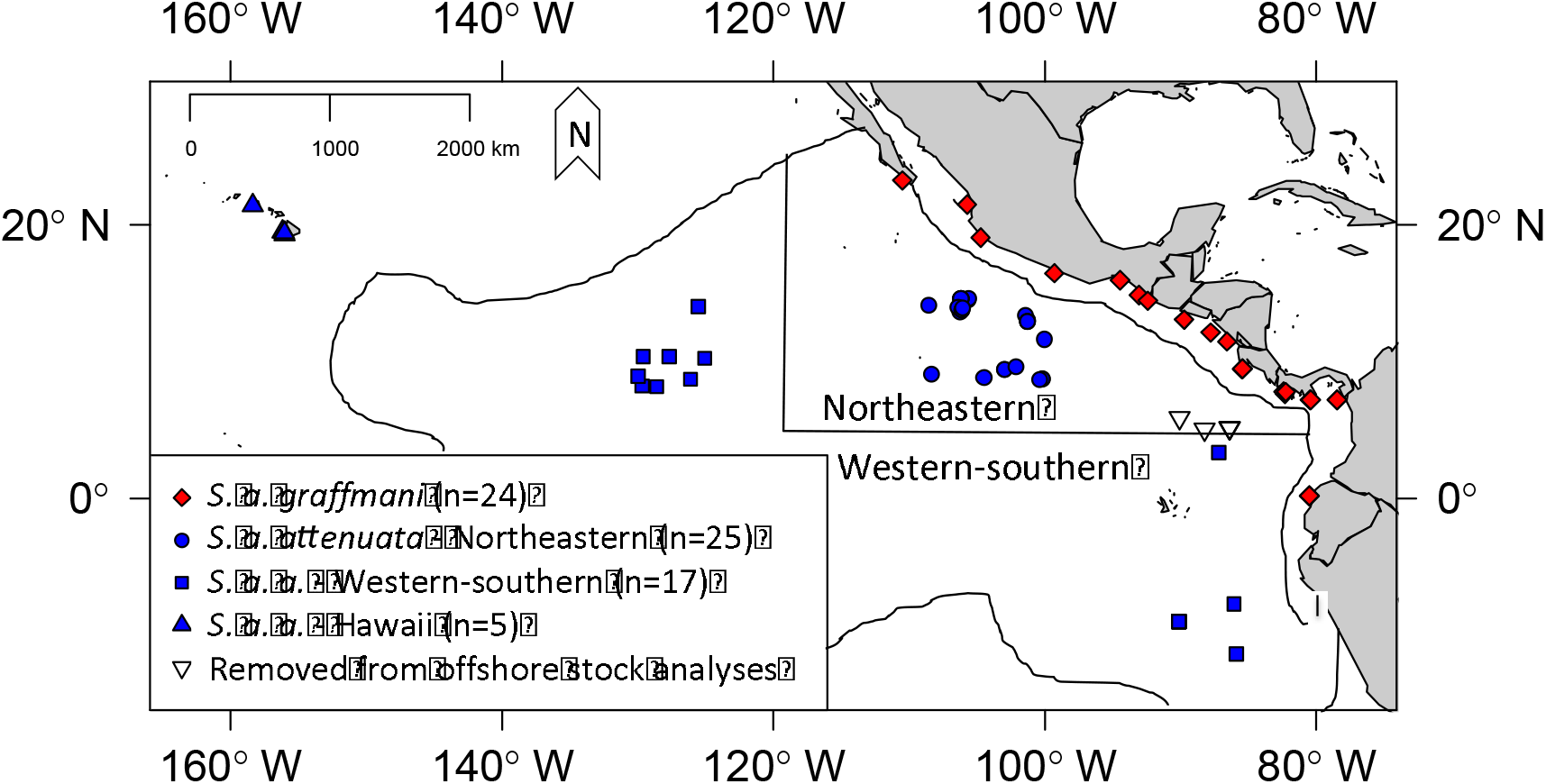
Sampling localities for spotted dolphins with ETP subspecies and stock boundaries based on Perrin *et al.* 1985. Coastal spotted dolphins (*S. a. graffmani*) are in red and offshore (*S a. attenuata*) are in blue. Blue circles indicate sampling locations for the northeastern stock of offshore spotted dolphins. Blue triangles indicate samples from Hawaii. Inverted triangles indicate southern offshore samples that were removed from analyses of offshore stocks because they were collected between 4°N and 6°N; these samples were included in subspecies-level analyses. Animals that represent the western substock were the group of blue squares west of 120°W and animals representing the southern sub-stock were the group of blue squares taken from south of the 5°N stock boundary. Samples sizes for mtDNA analyses presented in the legend.

Given the morphological differentiation between subspecies and recent evidence of nuclear DNA genetic differentiation, we assume the previous results from single mtDNA loci lacked power to resolve these close intraspecific relationships. In addition, studies of other cetacean species have shown mitochondrial genomes to be a useful tool for resolving intraspecific relationships when single mtDNA genes cannot (Archer *et al.*2013, Morin *et al.* 2010.

## Objectives

We used DNA capture array library enrichment and highly paralleled DNA sequencing to collect whole mitochondrial genome sequence data from 104 spinner and 76 spotted dolphins to test hypotheses of population genetic structure at multiple hierarchical levels in the eastern tropical Pacific Ocean. We performed analyses of whole mtDNA genomes (mitogenomes) and individual mtDNA genes to test observed levels of differentiation between recognized and proposed management stocks. We also tested for structure supporting the Tres Marias spinner dolphin and alternative stock boundaries in the offshore spotted dolphins. Although still only representing one locus, mitogenomes allow us to examine matrilineal population structure and contrast our findings with those found in previous studies using nuclear DNA (Escorza-Treviño *et al.* 2005; Andrews *et al.* 2013; Leslie and Morin 2016) to infer sex-biased dispersal.

## METHODS

### Sample Collection and DNA extraction

Skin samples used in this study were collected from free-ranging animals *via* biopsy dart (Lambertsen 1987) on research cruises or from dead specimens killed as bycatch in the tuna purse-seine fishery between 1982 and 2010 (104 spinner dolphins and 76 spotted dolphins Fig. 1, 2; Supplementary Material Tables S1, S2). On research cruises it is relatively common to see some fraction of spinner dolphins of alternate morphology (*i.e*., possibly different subspecies) within a school of dolphins comprised mostly of another morphotype/subspecies. For this reason, spinner dolphin samples collected from research cruises were assigned to a stock based on the external morphology of the majority of animals in the school, rather than the morphology of the individual sampled or the geographic location of the school. This method was preferable because: 1) only after observing the group (which could contain > 1,000 individuals) for some time could observers classify it to stock, 2) the external characters distinguishing subspecies are subtle, therefore researchers collecting biopsies from the bow of the research vessel could not confidently classify fast-swimming individuals in real time, and 3) the ranges of ETP spinner dolphin subspecies overlap making geography an unreliable predictor of stock identity. Some samples were used from areas where the eastern and whitebelly spinners are known to geographically overlap (see Figure 1). Spinner dolphin samples from Hawai‘i spanned the breadth of the main islands and also Midway Atoll.

Because there is little overlap of subspecies distribution in ETP pantropical spotted dolphins, geographic location of the sampling site was used to assign samples to subspecies and stocks. To avoid misassigned individuals near the borders of the NE and WS offshore stocks, we did not use samples collected between 4°N and 6°N east of 125°W. Hawai‘ian spotted dolphin samples were collected from the Kona Coast of Hawai‘i and O‘ahu.

Biopsy samples were stored in salt-saturated 20% DMSO, 70% ethanol, or frozen with no preservative. We extracted DNA using silica-based filter membranes (Qiagen, Valencia, CA) on an automated workstation (Perkin Elmer, Waltham, MA). DNA was quantified using Pico-Green fluorescence assays (Quant-it Kit, Invitrogen, Carlsbad, CA) and a Tecan Genios microplate reader (Tecan Group Ltd, Switzerland).

### Library Preparation and Sequencing

Next-generation sequencing libraries were generated as described by Hancock-Hanser *et al.* (2013), using unique 6bp and 7bp index sequences for each individual to allow up to 100 samples to be multiplexed. Multiplexed libraries were enriched for whole mitogenomes and 85 nuclear DNA loci using Sure Select DNA Capture Arrays (Agilent Technologies, Inc., Santa Clara, CA, USA) as described by Hancock-Hanser *et al.* (2013). Sequence data from the 85 nuclear loci were not used in this study. Target sequences for capture enrichment included the reference pantropical spotted dolphin mitochondrial genome (Genbank No. EU557096; Xiong *et al.* 2009) and a suite of 85 nuclear loci (not included in this study). Three identical arrays - designed with the eArray software package (Agilent Technologies, Inc., Santa Clara, CA, USA) - were used to capture a multiplexed mix of both species. Each array contained one replicate of the mitogenome probes at a probe interval of 15bp as well as 13 replicates of probes for the nuclear loci at a probe interval of 3bp. Each enriched library was then sequenced using 1X100bp Illumina HiSeq technology (two using Illumina HiSeq2000 and one using HiSeq2500).

### Mitogenome Assembly

Raw read data were filtered for quality (minimum quality score of 15) and demultiplexed by unique barcode. Consensus sequences for each sample were generated from mitogenome sequence reads using a custom pipeline (Dryad data repository doi:10.5061/dryad.cv35b) in R v2.15.0 (R Core Team, 2014). Reads were first mapped to the reference spotted dolphin sequence with the short-read alignment tool BWA (Li and Durbin, 2009). The mpileup module in SAMTOOLS (Li *et al.* 2009) was then used to convert the resulting BAM-format alignment file into a ‘‘pileup’’ text format, which was then parsed by custom R code to create the consensus sequence for each individual. The following rules were used in this process: A “N” was inserted at a position if the assembly had <3 reads, <5 reads where not all contained the same nucleotide, or >5 reads where no one nucleotide (*i.e*., A, C, G, T) was present in >70% of the reads. All mitogenome sequences were initially aligned with MAFFT using the automatic selection of an appropriate handling strategy (“auto”) and default parameters (Katoh *et al.* 2009) followed by a refinement of alignments by eye.

### Diversity Estimates and Population Structure Analyses

Two mitogenome data sets were created for each species. First, we partitioned each species’ mitogenome into fifteen loci (12 coding sequences, the control region and 2 rRNA genes). ND6 and tRNA loci were removed prior to analyses because they conform to different evolutionary models and ND6 falls on the opposite strand from the remaining genes (Duchene *et al.* 2011). Sequences were aligned to the pantropical spotted dolphin reference and locus start/stop positions were annotated in GENEIOUS v5.4 (Biomatters Limited) using the GENEIOUS alignment tool and the amino acid translation tool, respectively.

Second, we removed the control region because of high variation in this region and concatenated the remaining 14 regions to make the concatenated mitogenome sequences. The final sequence lengths for the concatenated data were 13,426bp and 13,425bp for spinner and spotted dolphins, respectively. An individual was removed entirely from analyses if it contained >10% missing data across the entire concatenated sequence.

For both data sets, we estimated haplotypic diversity (*h*, Nei 1987) and nucleotide diversity (*π*, Tajima 1983), and assigned individual genes and whole mitochondrial genome sequences to unique haplotypes using tools from the *strataG* package in R (v. 2.3.1; Archer *et al.*2016). Two pairwise estimates of population genetic structure, *F*_ST_ (Wright 1949) and *Φ*_ST_ (Excoffier *et al.* 1992), were also performed using the *strataG* package. The significance of each estimate was tested using 5000 non-parametric random permutations of the data matrix variables. For *Φ*_ST_, pairwise distances were calculated using the best substitution model as identified by Akaike’s Information Criterion in JModelTest version 2.1.4 (Posada 2008). Models were determined for individual gene regions and the entire concatenated dataset.

We performed a substitution rate test on each species’ mitogenome data set to determine if mutations had reached a point of saturation. For this test, we generated pairwise percent differentiation and plotted this against a Jukes and Cantor (1969) correction factor generated using MEGA 5.2.2 (Tamura *et al.* 2011). We chose this model because of its simplicity; if deviations were seen here then general saturation could be assumed.

Although mitochondrial loci are assumed to be under purifying selection (Stewart *et al.* 2008) we, nonetheless, tested spinner dolphin mitochondrial genes for evidence of positive selection using both Tajima’s *D* and Codon-based *Z*-Test as implemented in MEGA 5.2.2 (Tamura *et al.* 2011).

## RESULTS

Hancock-Hanser *et al.* (2013) present information on the success rate of the DNA capture method including summary statistics of the data analyzed in this paper. As it relates to our analyses, questions might arise about how using arrays designed from closely-related species affected our results. As presented in Tables 4 and 5 of Hancock-Hanser *et al.* (2013), spinner dolphin samples had slightly higher number of mtDNA reads per individual than spotted dolphin samples, despite use of the spotted dolphin mitogenome as the capture bait. The same pattern was found for the nuDNA capture – spinner dolphins had more reads per individual than spotted dolphins - despite all the baits being common bottlenose dolphin DNA sequence. We interpreted this consistency as an indication that inter-specific capture worked well and that any decrease in capture success (as evidenced in reads per individual for a given species) was more likely due to a combination of other factors (sample quality, multiplexing rate, sequencing technology, and/or variation in library preparation) rather than reduced capture due to inter-specific baits. The one area that might have been an issue for inter-specific capture was the hyper-variable section of the control region (see below).

### Spinner dolphins

We assembled 104 complete or nearly complete (<10% missing data) concatenated spinner dolphin mtDNA data sets (Genbank accession numbers in Supplementary Table S1). The hyper-variable section of the control region had consistently lower coverage in many individuals and was removed from the concatenated data set (Supplementary Table S1). Subspecies and regional sample sizes, summary statistics and genetic diversity measures are listed in Table 1. At the subspecies level, haplotypic diversities were high and nucleotide diversity was low (>0.9722, <0.0073, respectively). The substitution rate test did not show any signs of saturation. The best nucleotide substitution model estimated by JModelTest (Posada, 2008) was JC69 (Jukes and Cantor 1969) for each individual gene region and the entire concatenated data set. The results of *F*_ST_ and *Φ*_ST_ analyses of the mtDNA concatenated genes and *Φ*_ST_ of the individual gene regions for spinner dolphins are shown in Table 2. Due to space limitations, we only discuss *Φ*_ST_ for the partitioned gene region analyses.

**Table 1:** Summary statistics for ETP spinner (A) and spotted (B) dolphin mitogenome data. n_H_: number of haplotypes; PS: polymorphic sites; h: haplotype diversity; *π*: nucleotide diversity; %: percent of unique haplotypes.

| A. Spinner dolphins *Stenella longirostris* (n = 104) |  |  |  |  |  |  |
| --- | --- | --- | --- | --- | --- | --- |
| Subspecies/Stock | n:female/male/unk | n<sub>H</sub> | PS | h | $\pi$ | % |
| Central American<br>*S. l. centroamericana* | 9:4/4/1 | 8 | 648 | 0.9722 | 0.0057 | 0.7778 |
| eastern~<br>*S. l. orientalis* | 53:28/19/6 | 51 | 648 | 0.9985 | 0.0073 | 0.9245 |
| Putative Stocks |  |  |  |  |  |  |
| whitebelly<br>*S. l. longirostris* | 27:16/11/0 | 27 | 457 | 1 | 0.0043 | 1 |
| Tres Marias^~<br>*S. l. orientalis* | 21:8/10/3 | 20 | 373 | 0.9952 | 0.0078 | 0.9048 |
| Hawai'i<br>*S. l. longirostris* | 15:1/4/10 | 9 | 104 | 0.9921 | 0.0068 | 0.8260 |

| B. Spotted dolphins *Stenella attenuata* (n = 76) |  |  |  |  |  |  |
| --- | --- | --- | --- | --- | --- | --- |
| Subspecies | n:female/male/unk | n<sub>H</sub> | PS | h | $\pi$ | % |
| Coastal<br>*S. a. graffmani* | 24:11/13/0 | 16 | 234 | 0.9529 | 0.0162 | 0.5000 |
| ETP offshore<sup>§</sup><br>*S. a. attenuata* | 47:20/19/8 | 43 | 519 | 0.9804 | 0.0198 | 0.7222 |
| Offshore Stocks (*S. a. attenuata*) - Current and Putative^ |  |  |  |  |  |  |
| northeastern | 25:10/8/7 | 22 | 400 | 0.9867 | 0.0238 | 0.8000 |
| western-southern | 17:9/7/1 | 17 | 298 | 1 | 0.0096 | 1 |
| Offshore western^ | 8:7/1/0 | 8 | 191 | 1 | 0.0087 | 1 |
| Offshore southern^ | 9:2/6/1 | 9 | 253 | 1 | 0.0092 | 1 |
| Hawai'i | 5:1/3/1 | 3 | 36 | 0.7000 | 0.0244 | 0.4000 |

^ Stocks that are not recognized for management purposes. ~ The Tres Marias spinner samples are part of the eastern stratum. <sup>§</sup> Includes data for five samples that were omitted from stock comparisons because they were sampled too close to geographic stock boundaries.

**Table 2:** Pairwise divergence estimates for subspecies and stocks of spinner dolphins based on concatenated mitogenome data (*F*_ST_, *Φ*_ST_ and χ^2^) and d mitogenomic data (*Φ*_ST_ only). Light gray backgrounds for p<0.05; medium gray for p<0.01; darker gray backgrounds for p<0.001 (p-parentheses).

| Taxon 1 (n)<br>vs.<br>Taxon 2 (n) | Concatenated<br>mitogenome | | Partitioned mitogenome $\Phi_{ST}$ (p-value) | | | | | | | | | | | | | | |
| --- | --- | --- | --- | --- | --- | --- | --- | --- | --- | --- | --- | --- | --- | --- | --- | --- | --- |
| | $F_{ST}$<br>(p-value) | $\Phi_{ST}$<br>(p-value) | 12s<br>$n_H=24$ | 16s<br>$n_H=24$ | ATP6<br>$n_H=53$ | ATP8<br>$n_H=11$ | COI<br>$n_H=65$ | COII<br>$n_H=39$ | COIII<br>$n_H=47$ | CYTB<br>$n_H=61$ | CR<br>$n_H=50$ | ND1<br>$n_H=59$ | ND2<br>$n_H=53$ | ND3<br>$n_H=21$ | ND4<br>$n_H=56$ | ND4L<br>$n_H=22$ | ND5<br>$n_H=70$ |
| Central Amer. (9)<br>vs. eastern (53) | 0.0133<br>(0.034) | -0.0127<br>(0.5235) | -0.0120<br>(0.5265) | 0.0061<br>(0.4977) | -0.0076<br>(0.4001) | 0.0590<br>(0.0801) | -0.0276<br>(0.8640) | -0.0199<br>(0.6988) | -0.0158<br>(0.5766) | -0.0094<br>(0.4711) | 0.0017<br>(0.3983) | 0.0148<br>(0.2501) | -0.0260<br>(0.7376) | 0.0338<br>(0.1325) | -0.0287<br>(0.6950) | -0.0268<br>(0.7444) | -0.0139<br>(0.5368) |
| Central Amer. (9)<br>vs. whitebelly (27) | 0.0128<br>(0.056) | 0.0490<br>(0.0542) | -0.0165<br>(0.5882) | 0.0217<br>(0.1947) | 0.0311<br>(0.1277) | 0.1279<br>(0.0189) | 0.0351<br>(0.0903) | 0.0936<br>(0.0144) | 0.0086<br>(0.2995) | 0.0601<br>(0.0456) | 0.0555<br>(0.0412) | 0.0844<br>(0.0362) | 0.0113<br>(0.2833) | 0.1505<br>(0.0054) | 0.0870<br>(0.0464) | 0.0478<br>(0.0931) | 0.0273<br>(0.1203) |
| eastern (53) vs.<br>whitebelly (27) | 0.0007<br>(0.2867) | 0.0181<br>(0.0741) | 0.0307<br>(0.0414) | 0.0159<br>(0.0835) | 0.0051<br>(0.2421) | -0.0065<br>(0.5546) | 0.0264<br>(0.0288) | 0.0342<br>(0.0152) | -0.0020<br>(0.4501) | 0.0154<br>(0.1165) | 0.0270<br>(0.0059) | 0.0260<br>(0.0468) | 0.0104<br>(0.1687) | 0.0638<br>(0.0018) | 0.0464<br>(0.0422) | 0.0343<br>(0.0214) | 0.0026<br>(0.2859) |
| Tres Marias (21) vs.<br>Central Amer. (9) | 0.0155<br>(0.0914) | -0.0345<br>(0.7576) | 0.0113<br>(0.2921) | -0.0283<br>(0.7240) | -0.0436<br>(0.7284) | -0.0082<br>(0.2863) | -0.0393<br>(0.8636) | -0.0318<br>(0.7150) | -0.0301<br>(0.6752) | -0.0238<br>(0.5872) | -0.0102<br>(0.5328) | -0.0158<br>(0.5219) | -0.0451<br>(0.8698) | 0.0022<br>(0.4025) | -0.0558<br>(0.8900) | -0.0638<br>(0.9470) | -0.0383<br>(0.7888) |
| Tres Marias (21) vs.<br>eastern (32) | 0.0009<br>(0.4107) | -0.0116<br>(0.7084) | -0.0109<br>(0.6474) | -0.0217<br>(0.9462) | -0.0088<br>(0.5169) | 0.0019<br>(0.3119) | -0.0124<br>(0.7654) | -0.0182<br>(0.8772) | -0.0150<br>(0.8116) | -0.0062<br>(0.5291) | -0.0135<br>(0.8454) | -0.0117<br>(0.6898) | -0.0031<br>(0.4447) | -0.0206<br>(0.8894) | -0.0185<br>(0.7898) | -0.0049<br>(0.4887) | -0.0105<br>(0.6442) |
| Tres Marias (21) vs.<br>whitebelly (27) | 0.0024<br>(0.1934) | 0.0263<br>(0.0807) | 0.0421<br>(0.0643) | 0.0111<br>(0.1979) | 0.0086<br>(0.2423) | 0.0175<br>(0.2421) | 0.0323<br>(0.0519) | 0.0406<br>(0.0362) | -0.0005<br>(0.3907) | 0.0311<br>(0.0765) | 0.0413<br>(0.0168) | 0.0359<br>(0.0636) | 0.0243<br>(0.1087) | 0.0676<br>(0.0052) | 0.0859<br>(0.0789) | 0.0485<br>(0.0448) | 0.0124<br>(0.1807) |
| Hawaii (15) vs,<br>whitebelly (27) | 0.0456<br>(0.0001) | 0.1964<br>(0.0002) | 0.0236<br>(0.1667) | 0.3590<br>(0.0002) | 0.2560<br>(0.0002) | -0.0127<br>(0.6582) | 0.1964<br>(0.0002) | 0.1885<br>(0.0006) | 0.0154<br>(0.1363) | 0.1031<br>(0.0004) | 0.0771<br>(0.0036) | 0.0818<br>(0.0026) | 0.3302<br>(0.0002) | 0.4467<br>(0.0002) | 0.1324<br>(0.0002) | -0.0137<br>(0.2197) | 0.1858<br>(0.0002) |
| Hawaii (15) vs.<br>eastern (53) | 0.0449<br>(0.0001) | 0.1849<br>(0.0002) | 0.0428<br>(0.0605) | 0.3293<br>(0.0002) | 0.2268<br>(0.0002) | -0.0002<br>(0.4031) | 0.2061<br>(0.0002) | 0.2104<br>(0.0002) | 0.0338<br>(0.0625) | 0.1182<br>(0.0026) | 0.2050<br>(0.0002) | 0.1406<br>(0.0012) | 0.3090<br>(0.0002) | 0.3283<br>(0.0002) | 0.1339<br>(0.0025) | 0.0170<br>(0.1643) | 0.1494<br>(0.0007) |
| Hawaii (15) vs.<br>Central Amer. (9) | 0.0636<br>(0.0219) | 0.3284<br>(0.0002) | -0.0083<br>(0.4045) | 0.5265<br>(0.0002) | 0.3328<br>(0.0004) | 0.1600<br>(0.0631) | 0.3983<br>(0.0002) | 0.4280<br>(0.0002) | 0.1415<br>(0.0034) | 0.2474<br>(0.0002) | 0.2464<br>(0.0002) | 0.3091<br>(0.0002) | 0.4352<br>(0.0004) | 0.3863<br>(0.0002) | 0.2728<br>(0.0002) | 0.1597<br>(0.0701) | 0.2854<br>(0.0004) |
| Hawaii (15) vs.<br>Tres Marias (21) | 0.0487<br>(0.0004) | 0.2260<br>(0.0002) | 0.0796<br>(0.0478) | 0.3900<br>(0.0002) | 0.2552<br>(0.0002) | 0.0339<br>(0.2507) | 0.2576<br>(0.0002) | 0.2351<br>(0.0002) | 0.0703<br>(0.0272) | 0.1398<br>(0.0004) | 0.1533<br>(0.0002) | 0.1958<br>(0.0002) | 0.3454<br>(0.0002) | 0.3608<br>(0.0002) | 0.1667<br>(0.0004) | 0.0630<br>(0.0669) | 0.1828<br>(0.0002) |

At the subspecies level, the *Φ*_ST_ test showed no differentiation between Central American spinners and eastern spinner dolphin subspecies in either the concatenated or partitioned data sets. *F*_ST_ was significant in the concatenated data set (0.0133, *P =* 0.034). *Φ*_ST_ comparisons of the whitebelly form and coastal Central American subspecies showed nearly significant differentiation in the concatenated data set (*Φ*_ST_ = 0.0490; *P* = 0.0542) and seven individual gene regions. ND3 showed a significant difference at *P* = 0.0054, while all other significant comparisons between these strata were at *P* < 0.05 (Table 2).

We found no significant differences between the whitebelly and the eastern subspecies using the concatenated mitogenome data (*Φ*_ST_ = 0.0181; *P* = 0.0741). However, eight individual mitochondrial genes showed significant differentiation. All individual gene partitions in spinner dolphins were found to be under purifying selection using Tajima’s *D* tests for selection (Table S3) and Z-Test for positive selection using the Nei-Gojobori method (Nei and Gojobori 1986) (Table S6).

*Φ*_ST_ tests showed no differentiation between Tres Marias spinners and either ETP spinner dolphin subspecies in either the concatenated or partitioned data sets. Four individual gene regions were significantly different in the pairwise comparisons of Tres Marias and whitebelly spinner dolphins (*P* < 0.05; ND3 at *P* < 0.01).

All tests involving comparisons with Hawai‘ian spinner dolphins (*S. l. longirostris*) - using the concatenated data set - were highly significant. Four genes showed population structure (significant *Φ*_ST_) in all pairwise comparisons between Hawai‘i and ETP groups (*i.e*., Central America, Tres Marias, eastern, and whitebelly spinner), but not in any pairwise comparisons between these ETP groups: 16S, ATP6, ND2, and ND5. Because of the low abundance and geographic isolation of the Hawai‘ian population, we presume these genetic differences between Hawai‘i and the ETP groups resulted primarily from drift in the Hawai‘ian population, though some unique haplotypes in Hawai‘i also suggest sequence divergence between the subspecies. 16S had 24 haplotypes total, but only 4 haplotypes among all 15 Hawai‘ian samples. ATP6 had many more haplotypes in total (53), but again reduced diversity in Hawai‘i (5). One of these Hawai‘ian haplotypes was common among all ETP groups, and two were exclusive to Hawai‘i. The final two Hawai‘ian ATP6 haplotypes were shared with one ETP spinner dolphin each. ND2 also had 53 haplotypes total, but only 4 spread among the 15 Hawai‘ian samples. Twelve samples from Hawai‘i had two haplotypes that were not shared with ETP populations. One individual shared a haplotype with an eastern spinner dolphin, the other two haplotypes were single samples unique to Hawai‘i. Finally, ND5 had 70 total haplotypes, but only 5 among the Hawai‘ian samples – none of which were shared with ETP populations.

### Spotted dolphins

We assembled 76 complete or nearly complete (<10% missing data) spotted dolphin mitogenomes (Genbank accession numbers in Supplementary Table S2). Sample sizes, summary statistics and genetic diversity measures are listed in Table 1. At the level of subspecies, nucleotide diversity was higher in spotted dolphins (>0.0162) than spinner dolphins. Haplotypic diversity (*h*) is high in both species (>0.9529), but ETP spotted dolphins subspecies have slightly lower levels (0.9529 and 0.9804 for the coastal and offshore groups, respectively) than spinner dolphin subspecies (0.9722 and 0.9985) in this region. The coastal ETP subspecies for both spinner and pantropical spotted dolphins in the ETP show reduced *h* compared to their offshore ETP counterparts (Table 1). Similar to the spinner dolphin mitogenome data, the substitution rate test did not detect any signs of saturation, and JC69 was the best substitution model for all individual gene regions and the entire concatenated data set.

Results of *F*_ST_ and *Φ*_ST_ analyses of the mtDNA concatenated genes and *Φ*_ST_ of the individual gene regions for spotted dolphins are presented in Table 4. Similar to the spinner dolphins, our analyses at the subspecies level for spotted dolphins (coastal *vs*. offshore) show no significant differentiation using *Φ*_ST_ for the concatenated or partitioned data sets. *F*_ST_ was significant in the concatenated data set (0.0125, *P* = 0.0402).

**Table 3:** Pairwise divergence estimates (***F*_ST_**) for spinner and spotted dolphin subspecies, respectively, using all nuclear SNPs, and using only neutral SNPs. Light gray backgrounds for p<0.05; Medium gray for p<0.01; darker gray backgrounds for p<0.001.

| Spinner dolphins |  | All 51 SNPs | 42 Neutral SNPs |
| --- | --- | --- | --- |
| Taxon 1<br>n:female/male/unk | Taxon 2<br>n:female/male/unk | $F_{ST}$<br>(p-value) | $F_{ST}$<br>(p-value) |
| Central American<br>7:3/3/1 | eastern<br>28:15/7/6 | -0.0023<br>(0.4485) | -0.0066<br>(0.5148) |
| Central American<br>7:3/3/1 | whitebelly<br>21:12/9/0 | 0.0148<br>(0.2607) | 0.0082<br>(0.3216) |
| eastern<br>28:15/7/6 | whitebelly<br>21:12/9/0 | 0.0297<br>(0.0059) | 0.0282<br>(0.0099) |
| Spotted dolphins |  | All 36 SNPs | 25 Neutral SNPs |
| Offshore<br>13:6/6/1 | Coastal<br>12:5/7/0 | 0.1711<br>(0.001) | 0.1493<br>(0.0005) |

**Table 4:** Pairwise divergence estimates for subspecies and stocks of spotted dolphins using concatenated mitogenome data (*F*_ST_, *Φ*_ST_ and χ^2^) and d mitogenomic data (Φ_ST_ only). n_H_ listed below each gene name is the number of haplotypes for that gene. Light gray backgrounds for medium gray for p<0.01; darker gray backgrounds for p<0.001 (p-values in parentheses). “*NA*” indicates comparisons where *Φ*_ST_ could not ted because all individuals in both strata share the same haplotype. “*” are where one stratum was n<5.

| Taxon 1 (n) vs.<br>Taxon 2 (n) | Concatenated mitogenome | | Partitioned mitogenome $\Phi_{ST}$ (p-value) | | | | | | | | | | | | | | |
| --- | --- | --- | --- | --- | --- | --- | --- | --- | --- | --- | --- | --- | --- | --- | --- | --- | --- |
| | $F_{ST}$<br>(p-value) | $\Phi_{ST}$<br>(p-value) | 12S<br>$n_H=6$ | 16S<br>$n_H=7$ | ATP6<br>$n_H=20$ | ATP8<br>$n_H=5$ | COI<br>$n_H=21$ | COII<br>$n_H=13$ | COIII<br>$n_H=11$ | CYTB<br>$n_H=20$ | CR<br>$n_H=20$ | ND1<br>$n_H=15$ | ND2<br>$n_H=17$ | ND3<br>$n_H=10$ | ND4<br>$n_H=23$ | ND4L<br>$n_H=2$ | ND5<br>$n_H=29$ |
| Coastal (24) vs.<br>Offshore (52) | 0.0125<br>(0.0402) | -0.0091<br>(0.4961) | 0.0089<br>(0.2553) | -0.0316<br>(0.9932) | -0.0149<br>(0.6788) | -0.0169<br>(0.7536) | 0.0018<br>(0.3265) | -0.0133<br>(0.6182) | -0.0243<br>(0.8890) | -0.0143<br>(0.6067) | -0.0122<br>(0.7336) | -0.0099<br>(0.5357) | -0.0198<br>(0.7610) | -0.0222<br>(0.9006) | -0.0042<br>(0.4085) | 0.0041<br>(0.3023) | -0.0056<br>(0.4217) |
| Northeastern (25) vs.<br>western-southern (17) | 0.0045<br>(0.2099) | -0.0076<br>(0.4111) | -0.0014<br>(0.3779) | 0.0079<br>(0.2841) | -0.0156<br>(0.5332) | 0.0057<br>(0.2691) | -0.0194<br>(0.6164) | 0.0003<br>(0.3637) | -0.0193<br>(0.5067) | -0.0211<br>(0.5423) | -0.0091<br>(0.5552) | 0.0086<br>(0.2585) | -0.0021<br>(0.3473) | -0.0068<br>(0.4139) | 0.0039<br>(0.3077) | -0.0038<br>(0.3771) | -0.0187<br>(0.5574) |
| Coastal (24) vs.<br>northeastern (25) | 0.0302<br>(0.0002) | -0.0082<br>(0.4405) | 0.0032<br>(0.3375) | -0.0326<br>(0.8096) | -0.0204<br>(0.7070) | -0.0061<br>(0.4689) | -0.0007<br>(0.3651) | -0.0060<br>(0.4325) | -0.0271<br>(0.7540) | -0.0201<br>(0.6148) | -0.0041<br>(0.4653) | 0.0031<br>(0.3041) | -0.0119<br>(0.4797) | -0.0105<br>(0.4947) | 0.0055<br>(0.2923) | 0.0016<br>(0.3249) | -0.0125<br>(0.5309) |
| Coastal (24) vs.<br>western-southern (17) | 0.0144<br>(0.0884) | -0.0342<br>(0.8102) | -0.0186<br>(0.5621) | -0.0356<br>(0.6598) | -0.0297<br>(0.7402) | 0.0081<br>(0.3153) | -0.0313<br>(0.8118) | -0.0355<br>(0.8624) | -0.0449<br>(0.8950) | -0.0385<br>(0.8666) | -0.0024<br>(0.8060) | -0.0392<br>(0.9142) | -0.0335<br>(0.7224) | -0.0393<br>(0.9112) | -0.0311<br>(0.7582) | -0.0360<br>(0.7624) | -0.0285<br>(0.6812) |
| Offshore southern (9)<br>vs. offshore western (8) | 0.0771<br>(0.2249) | 0.1666<br>(0.0668) | -0.1717<br>(0.0781) | 0.2129<br>(0.0618) | 0.1167<br>(0.1039) | 0.1117<br>(0.0801) | 0.1229<br>(0.0743) | 0.1361<br>(0.0939) | 0.1816<br>(0.0767) | -0.0471<br>(0.4611) | 0.0683<br>(0.1177) | 0.1767<br>(0.0575) | 0.1771<br>(0.0743) | 0.1155<br>(0.1183) | 0.2148<br>(0.0394) | 0.1382<br>(0.1231) | 0.1895<br>(0.0529) |
| Northeastern (25) vs.<br>offshore western (8) | 0.0027<br>(0.4291) | 0.1135<br>(0.0517) | 0.0853<br>(0.0945) | 0.1848<br>(0.0352) | 0.0728<br>(0.1223) | 0.1128<br>(0.0252) | 0.0575<br>(0.1397) | 0.1164<br>(0.0504) | 0.1117<br>(0.0749) | 0.0064<br>(0.3259) | 0.0348<br>(0.1635) | 0.1525<br>(0.0372) | 0.1179<br>(0.0775) | 0.1142<br>(0.0697) | 0.1497<br>(0.0394) | 0.1309<br>(0.0689) | 0.0894<br>(0.0957) |
| Northeastern (25) vs.<br>offshore southern (9) | 0.0073<br>(0.3755) | -0.0400<br>(0.7446) | -0.0242<br>(0.5728) | -0.0537<br>(0.8008) | -0.0468<br>(0.8162) | -0.0691<br>(0.9552) | -0.0291<br>(0.6287) | -0.0387<br>(0.7512) | -0.0491<br>(0.7828) | -0.0509<br>(0.8168) | -0.0300<br>(0.7886) | -0.0379<br>(0.7394) | -0.0392<br>(0.7150) | -0.0551<br>(0.9540) | -0.0238<br>(0.5626) | -0.0694<br>(0.9756) | -0.0327<br>(0.6092) |
| Coastal (24) vs.<br>offshore southern (9) | 0.0255<br>(0.0762) | -0.0130<br>(0.4065) | -0.0323<br>(0.5874) | -0.0277<br>(0.4713) | -0.0279<br>(0.5721) | -0.0147<br>(0.4071) | -0.0122<br>(0.4423) | -0.0079<br>(0.3971) | -0.0227<br>(0.4611) | -0.0579<br>(0.8690) | -0.0418<br>(0.8558) | -0.0012<br>(0.3477) | -0.0419<br>(0.6714) | -0.0309<br>(0.5854) | 0.0051<br>(0.3209) | -0.0160<br>(0.4301) | -0.0030<br>(0.3453) |
| Coastal (24) vs.<br>offshore western (8) | 0.0049<br>(0.4321) | 0.0749<br>(0.1331) | 0.1368<br>(0.0559) | 0.1366<br>(0.0855) | 0.0594<br>(0.1583) | 0.1363<br>(0.0167) | 0.0751<br>(0.1281) | 0.0406<br>(0.1953) | 0.0769<br>(0.1535) | -0.0089<br>(0.3361) | 0.0488<br>(0.1359) | 0.0609<br>(0.1541) | 0.0901<br>(0.1101) | -0.0372<br>(0.2059) | 0.1067<br>(0.0881) | 0.0239<br>(0.2425) | 0.0841<br>(0.1269) |
| Hawaii (5) vs. Coastal<br>(24) | 0.1430<br>(0.0026) | 0.2773<br>(0.0208) | 0.4166<br>(0.0019) | 0.2176<br>(0.0572) | 0.2767<br>(0.0174) | -0.0502<br>(0.5687) | 0.4032<br>(0.0028) | 0.1687<br>(0.0762) | 0.2175<br>(0.0585) | -0.2859<br>(0.0254) | 0.2037*<br>(0.0326) | 0.3643<br>(0.0049) | 0.1575*<br>(0.0947) | 0.3085<br>(0.0042) | 0.2660<br>(0.0252) | 0.2811<br>(0.0202) | 0.2541<br>(0.0244) |
| Hawaii (5) vs. Offshore<br>(47) | 0.1181<br>(0.0006) | 0.1582<br>(0.0389) | 0.1806<br>(0.0422) | -0.1282<br>(0.1107) | 0.1882<br>(0.0352) | -0.0459<br>(0.6156) | -0.2138<br>(0.0124) | -0.0818<br>(0.1323) | 0.1361<br>(0.0632) | -0.0849<br>(0.1481) | 0.1485*<br>(0.0475) | 0.2598<br>(0.0082) | 0.1146*<br>(0.1449) | 0.2609<br>(0.0054) | 0.1239<br>(0.0593) | 0.1689<br>(0.0545) | 0.1303<br>(0.0517) |
| Hawaii (5) vs.<br>northeastern (25) | 0.0576<br>(0.2709) | 0.1308<br>(0.0645) | 0.1153<br>(0.0962) | 0.0809<br>(0.1793) | 0.1584<br>(0.0353) | -0.0198<br>(0.4695) | -0.1981<br>(0.0206) | 0.0638<br>(0.1739) | 0.1099<br>(0.1123) | 0.1478<br>(0.0714) | 0.1372*<br>(0.0567) | 0.2499<br>(0.0102) | 0.0869*<br>(0.1279) | 0.2751<br>(0.0051) | 0.0984<br>(0.1029) | 0.1446<br>(0.0843) | 0.0951<br>(0.1355) |
| Hawaii (5) vs.<br>western-southern (17) | 0.2474<br>(0.0702) | 0.2273<br>(0.0238) | 0.2942<br>(0.0284) | 0.2139<br>(0.0774) | 0.2670<br>(0.0297) | -0.0542<br>(0.8308) | 0.2583<br>(0.0244) | 0.1353<br>(0.1133) | -0.1992<br>(0.0547) | 0.2259<br>(0.0286) | 0.1704*<br>(0.0547) | 0.3089<br>(0.0196) | 0.1793*(<br>(0.1473) | 0.2751<br>(0.0234) | 0.1925<br>(0.0356) | 0.2673<br>(0.0342) | 0.2062<br>(0.0366) |
| Hawaii (5) vs.<br>offshore western (8) | 0.4958<br>(0.0732) | 0.4932<br>(0.0179) | 0.4558<br>(0.0318) | 0.5298<br>(0.0168) | 0.4984<br>(0.0148) | 0.0285<br>(0.3925) | 0.4640<br>(0.0119) | 0.4572<br>(0.0364) | 0.5036<br>(0.0352) | 0.0013<br>(0.4061) | 0.3598*<br>(0.0771) | 0.5523<br>(0.0039) | 0.4484*<br>(0.0328) | 0.4915<br>(0.0033) | 0.5093<br>(0.0114) | 0.1309<br>(0.0689) | .0924<br>(0.1393) |
| Hawaii (5) vs.<br>offshore southern (9) | 0.1509<br>(0.2207) | 0.1274<br>(0.1167) | 0.3012<br>(0.0268) | 0.0306<br>(0.2757) | 0.1750<br>(0.0718) | 0.0443<br>(0.1961) | 0.2126<br>(0.0202) | 0.0036<br>(0.4437) | 0.0161<br>(0.3045) | 0.1384<br>(0.0872) | -0.1260*<br>(0.0865) | 0.2138<br>(0.0178) | 0.0039*<br>(0.2641) | -0.0551<br>(0.0206) | 0.0808<br>(0.1389) | 0.1382<br>(0.1231) | 0.5038<br>(0.0198) |

Estimates of differentiation between the current management stocks within the offshore subspecies (NE and WS stocks) using the whole mitogenome data and individual mtDNA genes showed no differences. Using *Φ*_ST_, no significant differences were observed between the coastal subspecies and the NE offshore stock, however *F*_ST_ (0.0302) was highly significant at *P* = 0.0002 between theses management units. Similarly, *Φ*_ST_ was not significant for pairwise comparisons of the Coastal subspecies and WS offshore stock using the concatenated data or individual genes.

Within the WS offshore stock, we found nearly significant *Φ*_ST_ differences between the southern and western offshore regions using the concatenated mitogenome (0.1666; *P* = 0.0668). One individual mtDNA gene (ND4) had significant differentiation (p <0.05) and three others had nearly significant p-values (16S, ND1, ND5).

Comparing separate western and southern portions of the WS stock to other partitions using the mitogenome data set also yielded no significant *Φ*_ST_ estimates. Our comparison of the NE stock to the western portion of the WS stock, however, was nearly significant using the concatenated mitogenome (*Φ*_ST_ = 0.1135; *P* = 0.0517) and four individual mtDNA genes showed significant *Φ*_ST_ differences (p<0.05). Neither data set showed significant differences between the NE stock and the southern portion of the WS stock for either statistic.

Comparison of the coastal subspecies to just the southern portion of the WS stock resulted in no significant *F*_ST_ or *Φ*_ST_ difference in the concatenated data set or individual gene regions. Between the coastal subspecies and western offshore portion of the WS stock, however, one individual gene region (ATP8) showed significant differentiation (*P* < 0.05), and one (12S) showed nearly significant differentiation (*P* = 0.0559). Ideally we would have partitioned the coastal subspecies south of central Mexico into the population units described by Escorza-Triveño *et al.* (2005), but our smaller sample size prevented us from doing this.

Significant differentiation was detected between Hawai‘i and the coastal subspecies, and between Hawai‘i and offshore spotted dolphins, in *F*_ST_ and *Φ*_ST_ of the concatenated data set. As expected, given this result, significant differentiation was detected in many individual mtDNA genes (see Table 4). We also detected significant differences between Hawai‘i and the NE stock in four genes, but not for the concatenated mtDNA data set (although it was nearly significant for *Φ*_ST_ at *P* = 0.0645). Hawai‘i and the WS stock were significantly different in the concatentated data set using *Φ*_ST_, and in nine individual genes (*P* < 0.05).

Finally, we also tested hypotheses of differences between Hawai‘i and divided western and southern portions of the WS stock. Hawai‘i and the western portion were differentiated using the concatenated dataset (*Φ*_ST_: 0.4932; *P* = 0.0179). Ten individual genes showed differentiation between these two strata (see Table 4). Hawai‘i and the southern portion of the WS stock were not differentiated based on our concatenated data sets, but did show significant differentiation in five individual genes (*P*<0.05).

## Discussion

Spinner and spotted dolphins in the eastern tropical Pacific offer a unique opportunity to study genetic differentiation at multiple scales in species with strong intraspecific morphological differences. Recent divergence, high genetic diversity, large population sizes, and ongoing gene flow likely contribute to low detectability of genetic divergence (Galver 2002, Escorza-Treviño *et al.* 2005, Andrews *et al.* 2013, Taylor and Dizon 1996, Waples 1998). Using complete mitogenomes, we found some genetic support for endemic subspecies of spinner and spotted dolphins, although the strength of this support varies between markers (see Table 5). We did not find support, however, for the division of offshore stocks of spotted dolphins; nor did we find separation of the Tres Marias spinner dolphins as an independent population. In contrast, nuclear SNP analysis recovered these stock-level differences (Leslie and Morin 2016). The difference in our findings compared to those of Leslie and Morin (2016) could reflect the limitations of our mtDNA data or something biologically meaningful about the populations.

**Table 5:** Summary table of pairwise comparisons using mtDNA and nuDNA data sets (sample sizes in parentheses). In the mtDNA column, a “□” denotes significance of whole mtDNA based on at least one measure (see Tables 2-4), ‘ns’ = non-significant, “ ∼ ” = indicating possible structure with *P*-value between 0.05 and 0.1. “# Genes” is the number of significant mtDNA genes (“∼” + # for nearly significant genes). For the nuDNA column, a “□” denotes significance and ‘*NA*’ denotes not tested (Leslie & Morin 2016; *Escorza-Treviño *et al*. 2005).

| Spinner dolphins | Taxon 1 (n<sub>mt</sub>/n<sub>nuc</sub>) | Taxon 2 (n<sub>mt</sub>/n<sub>nuc</sub>) | mtDNA |  | nuDNA |
| --- | --- | --- | --- | --- | --- |
|  |  |  | Whole | # Genes |  |
| Test of endemic subspecies | Central American (9/9) | eastern (53/36) | □ | 0 | □ |
| Testing whitebelly intergrade | Central American (9/7) | whitebelly (27/15) | ~ | 7; ~1 | □ |
| Testing whitebelly intergrade | eastern (54/36) | whitebelly (27/15) | ~ | 8; ~1 | □ |
| Alternative stock hypotheses | Tres Marias (21/12) | Central American (9/9) | ~ | 0 | □ |
| “?” | Tres Marias (21/12) | Eastern (32/36) | ns | 0 | □ |
| “?” | Tres Marias (21/12) | Whitebelly (27/12) | ~ | 4; ~5 | □ |
| “?” | Hawaii (15/0) | Whitebelly (27/0) | □ | 11 | NA |
| “?” | Hawaii (15/0) | Eastern (32/0) | □ | 11; ~1 | NA |
| “?” | Hawaii (15/0) | Central American (9/0) | □ | 12; ~1 | NA |
| “?” | Hawaii (15/0) | Tres Marias (21/0) | □ | 13; ~1 | NA |

| Spotted dolphins | Taxon 1 (n<sub>mt</sub>/n<sub>nuc</sub>) | Taxon 2 (n<sub>mt</sub>/n<sub>nuc</sub>) | mtDNA |  | nuDNA |
| --- | --- | --- | --- | --- | --- |
|  |  |  | Whole | # Genes |  |
| Testing subspecies | Offshore (52/13) | Coastal (24/27) | □ | 0 | □* |
| Testing existing stocks | Offshore NE (25/15) | Off. western-southern (17/16) | ns | 0 | □ |
| Testing existing stocks | Offshore NE (25/15) | Coastal (24/27) | □ | 0 | □ |
| Testing existing stocks | Offshore WS (17/16) | Coastal (24/27) | ~ | 0 | □ |
| Alternative stock hypotheses | Offshore southern (9/0) | Offshore western (8/0) | ~ | 1; ~9 | NA |
| “?” | Offshore NE (25/0) | Offshore western (8/0) | ~ | 4; ~7 | NA |
| “?” | Offshore NE (25/0) | Offshore southern (9/0) | ns | 0 | NA |
| “?” | Offshore southern (9/0) | Coastal (24/0) | ~ | 0 | NA |
| “?” | Offshore western (8/0) | Coastal (24/0) | ns | 1; ~3 | NA |
| “?” | Hawaii (5/0) | Coastal (24/0) | □ | 10; ~4 | NA |
| “?” | Hawaii (5/0) | Offshore (52/0) | □ | 6; ~4 | NA |
| “?” | Hawaii (5/0) | Offshore NE (25/0) | ~ | 4; ~4 | NA |
| “?” | Hawaii (5/0) | Offshore WS (17/0) | □ | 9; ~3 | NA |
| “?” | Hawaii (5/0) | Offshore western (8/0) | □ | 10; ~2 | NA |
| “?” | Hawaii (5/0) | Offshore southern(9/0) | ns | 5; ~3 | NA |

First, we will discuss the individual comparisons for each species, then finish with an overall discussion comparing our findings with those of others.

### Spinner Dolphins

Traditional *F*_ST_ was very low as expected, but supported endemic subspecies distinction (Central American and eastern). We found non-significant results from *Φ*_ST_ – a metric that includes differences in nucleotide divergence. Thus we conclude that haplotypes within these two subspecies are very similar, but that haplotype frequencies are significantly different.

Nevertheless, our results provide evidence of genetic differentiation between the accepted ETP endemic subspecies concordant with morphology (Perrin *et al.* 1991) and results from Andrews *et al.* (2013) who used data from the nuclear Actin gene, and Leslie and Morin (2016) who used restriction-associated nuDNA sequencing. Differences in ecological, distributional, morphological, nuDNA, and now mtDNA data support the recognition of these distinct subspecies.

Breeding biology and movement patterns could also affect the patterns we see between the Central American and eastern spinner dolphins. In particular, assortative mating can decrease *N*_*e*_, which could serve to amplify signal of structure in the nuDNA genome. The eastern spinner dolphin is thought to have a polygynous mating system (Perrin and Mesnick, 2003). Perrin and Mesnick (2003) concluded that relatively few males are involved with mating, serving to reduce *N*_*e*_ and potentially increase genetic structure (Perrin and Mesnick, 2003). Conversely, however, a skewed breeding system might also increase dispersal, as adult male dominance might promote movements of juvenile males which then become established breeders outside their natal range. Unfortunately, very little is known about the movement patterns of individual dolphins in the ETP, and less is known about differences in movement based on sex. High site fidelity in males could also restrict male-mediated geneflow between groups and increase relative signal in nuDNA analyses.

*F*_*ST*_ and *Φ*_ST_ tests between the whitebelly spinner and the Central American endemic ETP subspecies revealed nearly significant differentiation (Table 5), indicating possible separation. Nuclear SNP data support differences between these two groups (Leslie and Morin 2016). In our mitogenome data set, every whitebelly sample had a unique haplotype. As a result, frequency-based measures of differentiation such as *F-*statistics were likely underestimated.

Using slightly different samples, Andrews *et al.* (2013) also found differentiation between Central American and whitebelly spinners using mtDNA genes (control region and cyt*b*). We recovered the same pattern for cyt*b* and several others (Table 2). Moreover, Andrews *et al.* (2013) included 10 samples of Central American spinners that had questionable subspecific identity (based on further investigation of the sample collection records at SWFSC by MSL). These samples were initially identified as Central American spinners, but the confidence in the identification was low and they should have been labeled as “unidentified”. Given the uncertainly, these samples could have been from eastern spinner dolphins. Removal of these samples reduced our representation of Central American spinners (*n*=9), which may have impacted our ability to detect intraspecific structure. However, the Central American subspecies has lower relative abundance, and therefore might be expected to show higher levels of structure due to drift.

Two biological explanations for the possible differentiation between Central American and whitebelly spinners in the mtDNA are isolation by distance and admixture between whitebellies and Hawai‘ian spinners. These are the two most geographically distant putative populations of ETP spinner dolphins; therefore, isolation by distance could contribute to population genetic structuring. Admixture between the whitebelly and Hawai‘ian spinners would bring novel haplotypes from the Gray’s subspecies (Hawai‘i) into the whitebellies resulting in genetic structure.

We found nearly significant *Φ*_ST_ between the whitebelly spinner and the eastern spinner using the concatenated mitogenome data. In addition, we also found significant differences between these strata in eight individual mtDNA genes. Leslie and Morin’s (2016) SNP analysis supported differentiation of these two groups. Andrews *et al.* (2013) inferred high migration rates between whitebelly and eastern spinner dolphins (30.1 migrants per generation from whitebelly to eastern and 57.9 migrants from eastern to whitebelly). Despite this high rate of migration, we detected evidence of differentiation.

As discussed, the statistical power to estimate levels of migration between very large populations with low relative sample sizes is weak (Waples 1998, Taylor *et al.* 2000). For this reason, we did not estimate levels of migration for these data. Andrews *et al.* (2013) did estimate migration in ETP spinner dolphins and found lower, but significantly different from zero, rates of migration per generation between populations of Gray’s (Hawai‘ian and other Pacific Island groups) spinners and the whitebelly spinners (3.22 migrants per generation into Gray’s and 1.6 into whitebelly spinners). The rate of migration into Gray’s spinner populations from the eastern population was estimated to be less than one (0.82), but significantly different from zero. Although this was not a major focus of our study, the differences we detected between the Hawaiian population and the ETP pelagic populations were higher than any comparisons within the ETP, supporting the hypothesis that this is an insular population or possibly subspecies.

Differences in breeding systems could help drive or maintain differentiation between eastern and whitebelly spinner dolphins (Perrin and Mesnick 2003). A polgynous system in eastern spinners could result in higher site-fidelity and lower male *Ne* – both of which would accentuate signal in nuDNA population structure. Alternatively, as discussed above, admixture between whitebelly and Hawai‘ian spinners could also result in novel whitebelly genotypes resulting in higher apparent population structure between whitebelly and eastern spinners.

### Alternative spinner dolphin stocks

We found no support for a Tres Marias population that differs from the eastern or Central American subspecies (*e.g.,* Perryman and Westlake 1998) using the concatenated or individual mitochondrial gene data sets. Given the weak genetic differences we found between the accepted endemic subspecies with much more marked morphological differences, this result may not be surprising. We found statistically significant differences in four individual mtDNA genes when comparing the Tres Marias group to the whitebelly spinners and several nearly significant genes. We do not feel confident making taxonomic recommendations for the “Tres Marias” spinners based on these analyses. Additional studies should approach this question using larger sample sets and additional data.

### Spotted dolphins

Spotted dolphin mitogenomes have lower haplotypic diversity but higher nucleotide diversity than spinner dolphins, despite extremely high historical population sizes in the former. The two main reasons for lower haplotypic diversity could be a recent and/or prolonged population bottleneck, such as the decrease caused by mortalities in the tuna purse-siene fishery, or an extremely matrifocal social structure (Hoelzel *et al.* 2007). Although matrifocal social structure is known in several species of odontocetes (*e.g.,* killer whales and sperm whales), it is not a known characteristic of spotted dolphins, and thus is an unlikely cause of low genetic diversity.

Similar to our findings for spinner dolphins, traditional *F*_ST_ calculated for the spotted dolphin mitogenome data set supports differentiation of the offshore *S. a. attenuata* and the endemic coastal *S. a. graffmani* subspecies, whereas *Φ*_ST_ failed to indicate any difference - either for the entire genome or within any single gene. Our results show the NE stock being strongly differentiated from the coastal subspecies (based on allele frequency alone), counter to the results found by Escorza-Treviño *et al.* (2005) showing connection between the NE stock and the coastal subspecies based on seven microsatellite loci. In that study, the authors inferred that there was a strong connection between the coastal and offshore subspecies in northern Mexico. The differences between our results and those of Escorza-Treviño *et al.* (2005) could be due to sampling; the previous study included more samples from the northern portion of the coastal spotted dolphin range than we did. Additionally, the differences could be attributed to the unique evolutionary patterns of the different markers examined in Escorza-Treviño *et al.* (2005) (*i.e.,* microsatellites) vs. the mitogenomes used in our study.

### Spotted dolphin stocks

A main objective of this work was to test for difference between existing (NE, WS, and Coastal) and proposed (independent W and S) management stocks. Using the whole mtDNA genome data set, we found no evidence for differentiation between the two current stocks (NE and WS). This could be because the two stocks are genetically connected or because our data lack power to detect differentiation at this fine scale. The concatenated mtDNA indicated weak evidenced for splitting up the current WS stock - a high *Φ*_ST_ value (0.1666) and nearly significant (*P* = 0.0668). Similarly, we detected nearly significant differences between the NE stock and the western group of the WS stock using the concatenated mtDNA genome. Four mtDNA loci had significant *Φ*_ST_ estimates for this partition. The NE and the offshore southern group were not significantly different in any test, suggesting that the distributional hiatus at 5**°** north is not a barrier to gene flow. We cannot say with any certainty if this is the case, however, because of the low sample size for the southern portion of WS stock (n=9); a larger sample size is necessary to convincingly investigate this hiatus. Overall, the whole concatenated mtDNA genome was not as useful as anticipated for delimiting stock structure, possibly because it introduced more variation (*via* novel haplotypes) into an already highly variable system. Whole mtDNA genomes have been useful for clarifying subspecific boundaries where information in single mtDNA genes has shown low variability (Archer *et al.*2013, Morin *et al.* 2010), including in this paper, but testing population-level boundaries in highly abundant cetaceans using mtDNA genomes may be less feasible.

Leslie and Morin (2016) found divergence between the offshore and coastal spotted dolphin subspecies, but did not include data from individuals from the NE offshore stock of spotted dolphins. Therefore, this comparison includes animals from the most geographically separate portions of the offshore (WS) and coastal subspecies range. Additional nuclear data from the NE stock are needed to determine whether proximate populations of these two subspecies are also as genetically divergent.

### Overall Patterns Observed

Despite the increase in data over other mtDNA studies, it is likely that whole mitogenomes still do not provide enough statistical power to detect differences given the recent divergence, continued low-level interbreeding, and/or high diversity and historical abundance. We collected whole mitogenomic data to help resolve close population relationships. However, one risk of adding more data in this situation – with populations with high genetic diversity – is that haplotype discovery may not plateau within a sample set. In other words, more unique haplotypes are added thereby increasing the difficulty of characterizing haplotype frequencies among and between populations. Sequencing additional samples could help rectify this issue.

Similarly, there are also limitations to using frequency-based *F*-statistics. *F*_ST_ is good at detecting frequency differences that indicate genetic structure in cases where haplotypes are similar within populations and different between populations (such as those that would result via drift in small populations). However, when haplotype diversity is high within and among populations, very large sample sizes are needed to characterize haplotype frequencies to detect differences using *F*_ST_. In this situation, *F*_ST_ point values will be underestimated. Moreover, sampling effects can become important drivers of *F*_ST_ beyond the base frequency of alleles present and result in false positive results. Our initial hypothesis was that sequencing more of the mitogenome would result in more shared differences within populations which would be reflected in both the frequency-based statistics; this was not the case.

Instead, we found significant differences between subspecies of both spinner and spotted dolphins using *F*_ST_, but not *Φ*_ST_. *F*_ST_ and *Φ*_ST_ provide slightly different perspectives on population differentiation and we believe it is important to present both measures. Our results show inconsistencies between these two metrics, which does not necessarily mean analytical problems or inaccuracies, but reflects something interesting about our data. *F*_ST_ tests for population differentiation are based on allele (or haplotype) frequencies and do not provide direct insights into levels of molecular divergence (Weir and Cockerham, 1984, Excoffier *et al.* 1992, Meirmans and Hedrick 2011). *Φ*_ST_ estimates capture more information regarding the differentiation due to sequence divergence (or nucleotide diversity) in addition to differences in haplotype frequencies. The high heterozygosity issues mentioned above can still impact *Φ*_ST_. Although we chose to focus the bulk of the discussion on *Φ*_ST_, we do report statistically significant measures of *F*_ST_ and briefly compare and contrast the two metrics. *Φ*_ST_ may be more indicative of older, long-term processes, whereas *F*_ST_ can show recent differences among populations, indicating another reason it is important to report both metrics. In addition, given that the test for significance is determined by an arbitrary cut-off (*P* = 0.05), we also present results that are “nearly significant”. Combined with given the difficulty of distinguishing these groups in previous works, we felt it important not to focus too intensely on the arbitrary cut-off, but rather overall patterns of indicators.

Moreover, our sample sizes were low in some partitions (n=7). This could result in the allele frequencies of populations being under-characterized, which could skew results in over- or under-classification. Efforts should be made to collect more samples for future studies.

Alternatively, non-significant results could occur with low levels of geneflow between strata that are demographically independent populations (Avise 1995; Taylor and Dizon 1996). Because management decisions rely on them, results must be interpreted within the context of all available information and with recognition of the caveats of the data used to generate them.

The discordance we observed between the mitogenome results and those using nuDNA data (Leslie and Morin 2016) could also reflect biological factors. One possibility is female-mediated exchange diluting the signal of structure in mtDNA or male site-fidelity increasing structure in the mtDNA. Although there is some evidence from radio tagging studies that spinner and spotted dolphins can move relatively large distances (Perrin *et al.* 1979), a thorough investigation into the differences between sexes is lacking. At least for spinner dolphins it is likely that the polygynous breeding system described by Perrin and Mesnick (2003) would contribute to increased signal of structure in the nuclear genome.

### Drift in mtDNA loci as indicated by comparisons with Hawai‘i

Because of the greater divergence observed between Hawai‘ian and ETP populations of these two dolphins, we thought it would be informative to highlight genes showing structure (Hawai‘i vs. ETP), likely due to neutral drift acting on a small insular population, that might be useful for studying other Hawai‘ian populations of cetacean species. Four genes (16S, ATP6, ND2, and ND5) showed population structure (significant *Φ*_ST_) in all pairwise comparisons between Hawai‘i and ETP spinner dolphin groups (*i.e*., Central America, Tres Marias, eastern, and whitebelly spinner), but not in any pairwise comparisons between these ETP groups. All of the mtDNA regions with significant *Φ*_ST_ were found to be under purifying selection (negative Tajima’s *D* - Table S3; and non-significant Z-tests – Table S4) indicating that the within-mitogenome differences are accumulating by neutral drift rather than via positive selection in ETP spinner dolphins. Significant differences between ETP groups and the Hawai‘ian insular population of spotted dolphins were found in all but five of the mtDNA genes. We note however that the low sample sizes for Hawai‘ian spotted dolphins may explain some of the non-significant differences observed with respect to ETP stocks.

### Positive Selection in ETP Spinner Dolphin mtDNA

Selection should affect linked loci equally; however, selection can act on individual mtDNA genes, such as in the case of cytochrome *b* in Antarctic killer whales (Foote *et al.* 2010). We tested for positive selection in spinner dolphin mitochondrial genes and found none. We did not test for positive selection in spotted dolphins because there were no individual mtDNA genes that supported differentiation between the two ETP subspecies.

## Conclusions

Defining population genetic structure is challenging for species with large historical population sizes and high mobility. These populations may retain high genetic variation even as abundance becomes relatively low, which could obscure signals of genetic structure used to designate stock boundaries for estimating population abundance and setting stock-specific mortality limits. Ultimately, without information on structure, populations could be under-classified and unique evolutionary units and populations could go extinct as we may fail to take appropriate conservation action. Alternatively, there is a cost to managing populations as separate when there is no biological basis to do so. Such errors can have economic, social, and political consequences resulting from unnecessary restrictions on human activity. Furthermore, a consistent pattern of these errors will “stiffen the resolve of skeptics and make it difficult to accomplish sound resource management in the future” (Waples 1998).

This unique system of two delphinids, with available samples collected *in situ* from remote offshore environments encompassing extensive geographic and morphological variation, was used to test for population genetic structure at multiple hierarchical levels in species with high historical abundance and high intra-specific morphological variation. Our results show a complex pattern of genetic structure in the two different data sets for each species. Although complex, we believe the structure observed in our results is biologically meaningful. Given the aforementioned difficulties with detecting structure using genetic techniques in this system – and the supporting morphometric results - even subtle signatures of structure are significant findings. The mitogenome data show support for the endemic ETP spinner and spotted dolphin subspecies.

We found very little support for the division of offshore stocks of spotted dolphins and no support for the unique form of Tres Marias spinner dolphins as compared to the eastern or Central American subspecies. This is not to say that these biological entities do not exist, just that our mtDNA data do not support them or may not have sufficient power to detect the subtle genetic differences between them. Further, we recommend the collection and analysis of additional samples from the Central American subspecies to compare to existing offshore subspecies samples collected from fisheries bycatch and research cruises. In addition, we recommend additional studies of population structure that incorporate environmental variables as potential population boundaries in this area. Finally, placing these populations within a global phylogeographic context will help provide a better context for our results by fully characterizing intraspecific diversity and establishing the evolutionary process that led to ETP endemism.

## Acknowledgements

MSL was supported by a National Science Foundation (NSF) Graduate Research Fellowship, a NSF Integrative Graduate Education and Research Traineeship Fellowship, the Ralph A. Lewin Graduate Fellowship in Marine Biology at Scripps Institution of Oceanography, the Lerner-Grey Memorial Foundation at the American Museum of Natural History, and the Edna Bailey Sussman Foundation. We would especially like to thank Dr. William F. Perrin who provided guidance and support throughout the development and completion of this study. Danielle Davila, Brittany Hancock-Hanser, Gabriela Serra-Valente, Victoria Pease, Kelly Robertson, Dr. Louella Dolar, Nicole Beaulieu and Morgane Lauf for help in the laboratory and tissue archive. Dr. Karen Martien provided a thoughtful critique of this manuscript. Al Jackson provided copies of original field datasheets. We thank Dr. Robin Baird of Cascadia Research Collective and Dr. Erin Oleson of the NOAA Pacific Islands Science Center for permission to use Hawai‘ian spinner dolphin samples. The Scripps Research Institute provided assistance with sequencing, especially Drs. Steve Head, Lana Shaffer and John Shimashita. Support for this research was also provided from Drs. Lisa Ballance and Barbara Taylor of the Marine Mammal and Turtle Division of SWFSC. We are grateful to William Perrin, Karen Martien, Kim Andrews, Patricia Rosel and three anonymous reviewers for their careful and constructive reviews of the manuscript.

